# Establishment Process of a Magic Trait Allele subject to Both Divergent Selection and Assortative Mating

**DOI:** 10.1101/869198

**Authors:** T. Sakamoto, H. Innan

## Abstract

Sexual selection and divergent selection are among the major driving forces of reproductive isolation, which could eventually result in speciation. A magic trait is defined such that a single trait is subject to both divergent selection and sexual selection through phenotype-based assortative mating. We are here interested in the evolutionary behavior of alleles in a genetic locus responsible for a magic trait. We assume that, in a pair of homogeneous subpopulations, a mutant allele arises at the magic trait locus, and theoretically obtain the probability that the new allele establishes in the population. We also consider the trajectory of allele frequency along the establishment. Divergent selection simply favors the new allele to fix where it is beneficial, whereas assortative mating works against rare alleles. It is theoretically demonstrated that the fate of the new allele is determined by the relative contributions of the two selective forces, divergent selection and assortative mating, when the allele is rare so that the two selective forces counteract. We also show that random genetic drift also plays an important role. The theoretical results would contribute to improve our understanding of how natural selection initiates speciation.

---

Speciation occurs when reproductive isolation establishes between different populations. Sexual selection is one of the major forces driving reproductive isolation (Coyne and Orr 2004). Speciation driven by sexual selection could occur when phenotypic difference is involved in mate choice. Several theoretical models indicated that sexual selection alone can lead speciation even in the face of gene flow (Wu 1985; Turner and Burrows 1995; Higashi *et al.* 1999; Takimoto *et al.* 2000), but these results largely rely on their assumptions such as ample genetic variation, symmetric distribution of female preference or strong female choice (Arnegard and Kondrashov 2004; Gavrilets 2004), and empirically not well supported yet as reviewed in Ritchie (2007). At the moment, it is considered that speciation by sexual selection alone is difficult to occur (Gavrilets 2004). This is largely because diversity in female preference is difficult to maintain, rather disappears by genetic drift as female preference is not directly subject to selection (i.e., selection works through male phenotype). Ritchie (2007) pointed out that sexual selection should work efficiently together with niche specialization or local adaptation.

Synergy between local adaptation and assortative mating can be a powerful driver of speciation (Gavrilets 2004). Establishment of a locally adaptive mutation could lead stable genetic divergence between local populations in different environments, even in the face of gene flow between them. If there is another locus that is involved in sexual selection, it also reduces gene flow between populations. This effect is particularly strong when the locus is genetically linked to the target locus of local adaptation and an extreme case is that a single locus is pleiotropically subject to both divergent selection and sexual selection. Models that handle sexual selection at a locus under divergent selection are called “magic trait” models (Gavrilets 2004; Servedio *et al.* 2011). Previous theoretical studies revealed that the magic trait models are one of the easiest scenarios of speciation with gene flow (see Gavrilets 2004; Kopp *et al.* 2018, for reviews). There are many potential examples of a magic trait in nature (Maan and Seehausen 2011; Servedio *et al.* 2011), suggesting importance of the establishment process of a magic trait for speciation. In the present article, we specifically focus on a case of a single magic trait locus, which produces a phenotype difference and undergoes similarity-based mating (i.e., females prefer males with similar phenotypes).

Many theoretical models have considered a magic trait that is subject to both natural selection and similarity-based assortative mating (Dieckmann and Doebeli 1999; Matessi *et al.* 2001; Doebeli and Dieckmann 2003; Kirkpatrick and Nuismer 2004; Bürger and Schneider 2006; Otto *et al.* 2008; Pennings *et al.* 2008; Thibert-Plante and Gavrilets 2013; Rettelbach *et al.* 2013; Servedio and Bürger 2015; Cotto and Servedio 2017) (see Kirkpatrick and Ravigné 2002; Gavrilets 2004; Weissing *et al.* 2011; Servedio and Boughman 2017; Kopp *et al.* 2018, for review). However, these models often consider too complicated scenarios that, for example, a magic trait contributes to speciation in interaction with other traits or other selective forces, making it difficult to understand the relative contribution of the magic trait.

So far, theoretical arguments on the evolutionary dynamics of a magic trait have been only based on limited studies in an infinite population (Slatkin 1982; Kisdi and Priklopil 2011). Slatkin (1982) considered a haploid infinite-size island population connected to a stable continent population, between which migration is allowed. The model involves a single magic trait locus, at which two alternative alleles are considered: One allele is preferred in the island subpopulation, and the other allele is fixed in the continent subpopulation. In addition, assortative mating is incorporated by assuming that mating pairs of haploid individuals with different alleles produce fewer offsprings than pairs with the same alleles. Although this mode of selection is categorized into fecundity selection, it is mathematically identical to assortative mating, where females dislike to mate with males having different alleles. Slatkin (1982) analytically showed that successful invasion of a new allele requires a larger selection coefficient of adaptive selection than the sum of the migration rate and strength of assortative mating, both of which have an effect against the invasion of new alleles. Additionally, the author derived the critical migration rate, over which polymorphic state is unstable so that new alleles likely become extinct. Recently, Kisdi and Priklopil (2011) explored the evolutionary branching of a magic trait in an infinite population, although this model is not very relevant to our interest because the magic trait is quantitative determined by multiple genes. While the analysis assuming an infinite population gives a great deal of insight, it is also important to consider this process in a finite population with the stochasticity of random genetic drift. To our best knowledge, there has been no research which explored analytically the establishment process of a magic trait in stochastic two-population models, while some theoretical results are available in one-population models (Yamamichi and Sasaki 2013; Newberry *et al.* 2016).

To understand the effect of random genetic drift, we here consider a two-population model with bidirectional migration and derive the establishment probability of an allele which is pleiotropically subject to both divergent selection and assortative mating. Our model can arbitrarily set the sizes of the two populations, so that Slatkin (1982)’s situation (with the assumption of infinite population size) can be a special case of our model. We consider haploid and diploid models, and obtain theoretical expressions of the establishment probability and the trajectory of allele frequency along the establishment. Our derivation of the establishment probability largely relies on the approximation method developed by Yeaman and Otto (2011). Yeaman and Otto (2011) considered divergent selection alone in a two-population model and derived an approximate formula for the establishment probability of a locally adaptive allele (for theoretical works on this model of selection, see also Pollak 1966; Barton 1987; Tomasini and Peischl 2018; Sakamoto and Innan 2019). The authors developed a heuristic method, which essentially focuses on the process in the adaptive subpopulation, rather than considering the dual process in the adaptive and maladaptive subpopulations. This is because the establishment probability is largely determined by how the focal allele increases in frequency when it invades into the adaptive population, and the process in the maladaptive subpopulation does not play a crucial role. The authors showed that Kimura’s formula (Kimura 1962) well approximates the establishment probability when an adaptive allele arises in the adapted subpopulation if the selection coefficient is replaced by the leading eigenvalue of transition matrix of deterministic dynamics. In this work, following the idea of Yeaman and Otto (2011), we successfully derive the establishment probability in haploid and diploid models, not only when an adaptive allele arises in the adapted subpopulation but also when an adaptive allele arises in the maladapted subpopulation.

## MODEL

We use a discrete-generation two-population model, between which bidirectional migration is allowed, and we consider both haploid and diploid cases. The assumptions and model settings shared by the haploid and diploid models are described here. The sizes of subpopulations I and II are assumed to be *N*_1_ and *N*_2_, respectively. On average, *N*_1_*m*_1_ = *N*_2_*m*_2_ individuals are exchanged per generation where *m*_1_ and *m*_2_ are backward migration rate of subpopulation I and II, respectively. There are two alleles, A and a, on which selection works. Allele A is favored in subpopulation I, and disfavored in subpopulation II. Assuming no recurrent mutation between alleles A and a, we focus on the behavior of allele frequencies to obtain the establishment probability of allele A forward in time. Life cycle is assumed to be in the order of selection, mating and migration in each generation. The major difference between the haploid and diploid models is in how selection works. We assume that the fitness of allele a is 1, and the fitness of allele A is given depending on the model, as will be explained below.

### Haploid Model

In the haploid model, we simply assume that the fitness of allele A is 1 + *s*_1_ > 1 in subpopulation I, but 1 + *s*_2_ < 1 in subpopulation II. Let *p*_*i*_ be the allele frequency of allele A in subpopulation *i*. Then, the expected frequency of allele A after the selection event is

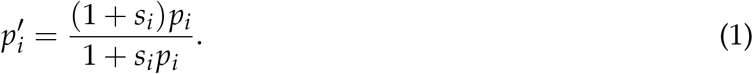

In the mating event, reproductive opportunity of a female is independent of her genotype. Assortative mating occurs such that a female avoids mating with male who has a different allele from hers with probability *α*. Then, the expected frequency of allele A after the mating event is given by

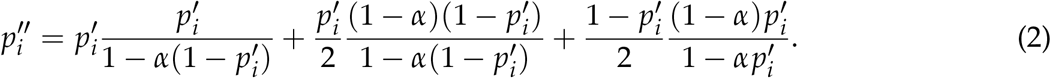

We assume that individuals that constitute the next generation are randomly produced by binomial sampling with 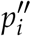. After this mating event, *N*_1_*m*_1_ = *N*_2_*m*_2_ individuals are exchanged between subpopulations in the migration event.

### Diploid Model

In the diploid model, we consider the dominance effect on the phenotype, based on which assortative mating works. We assume allele A has a dominance coefficient, *h*, which means that if genotype AA and genotype aa have trait values *P*_*AA*_ and *P*_*aa*_, respectively, the trait value of genotype Aa is given by *hP*_*AA*_ + (1 − *h*)*P*_*aa*_. We put *p*_*i*_, *q*_*i*_ and *r*_*i*_ as the genotype frequency of AA, Aa and aa, respectively, in subpopulation *i*.

In the selection event, we assume the fitness of genotype AA, Aa and aa are 1 + *s*_1_, 1 + *hs*_1_ and 1 in subpopulation I, and 1 + *s*_2_, 1 + *hs*_2_ and 1 in subpopulation II (*s*_1_ > 0 and *s*_2_ < 0). Then, the expectation of frequencies of AA, Aa after the selection event are given by

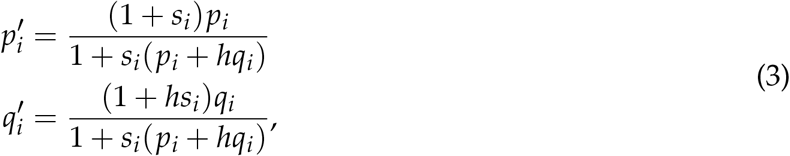

respectively, and the expected frequency of aa is given by 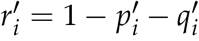.

In the mating event, a female avoids mating with a male who has a different trait from hers. To incorporate the effect of dominance, we assume that a female avoids mating proportional to the trait difference, and that genotype AA avoids mating with genotype aa with probability *α*. Then, the expected frequencies of genotype AA and aa after the mating event are

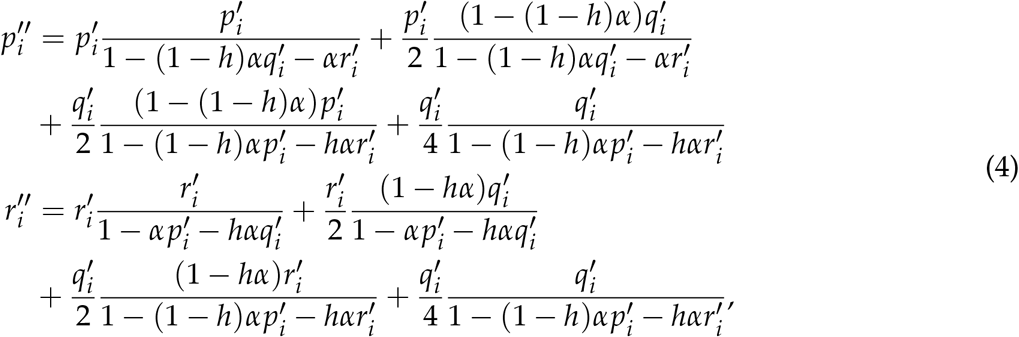

respectively, and the expected frequency of Aa are given by 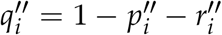. According to these probabilities 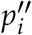 and 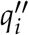, individuals for the next generation is produced by multinomial sampling. After that, *N*_1_*m*_1_ = *N*_2_*m*_2_ individuals are exchanged between subpopulations (migration event).

## RESULT

### Establishment Probability in the Haploid Model

The initial state is that all individuals have allele a in both subpopulations. First, we derive the establishment probability of allele A which arises in subpopulation I with initial frequency 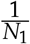. We assume *m*_*i*_ ≪ |*s_i_*|, *α* to ensure the stable maintenance of divergence (see below). By assuming that |*s_i_*|, *α*, *m_i_* ≪ 1 and ignoring the second order of these parameters, the expected changes in allele frequency in one generation are given by

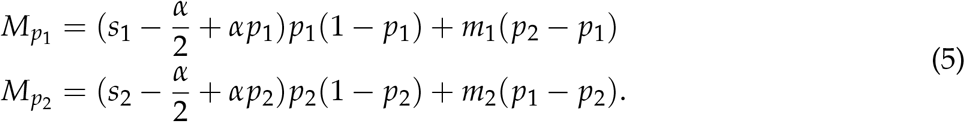

The change of allele frequencies, *p*_1_ and *p*_2_, can be well described by a two-dimensional diffusion equation, which is unfortunately very difficult to solve. Alternatively, we use the heuristic approach developed by Yeaman and Otto (2011), which stemmed from their demonstration that the establishment probability of an allele which arises in the adapted subpopulation can be approximated by the establishment probability of an allele under directional selection if the selection intensity is replaced by the leading eigenvalue of transition matrix of deterministic process. This means that the leading eigenvalue is considered to represent the ‘effective’ strength of natural selection on allele A which arises in subpopulation I.

The deterministic process considering divergent selection and migration (but not assortative mating) is described as

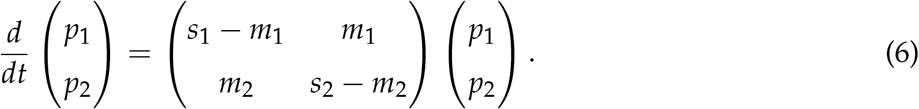

The leading eigenvalue of the transition matrix in Equation 6, *λ*, is given by

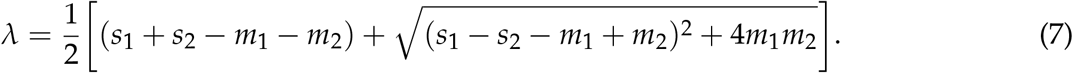

Following Yeaman and Otto (2011), the expected change in allele frequency in one generation is approximated in the one-population system:

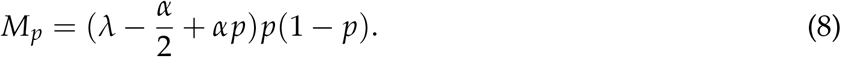

Then, the establishment probability of allele A which newly arises in subpopulation I, *u*_1_, is given along Kimura’s formula (Kimura 1962) as

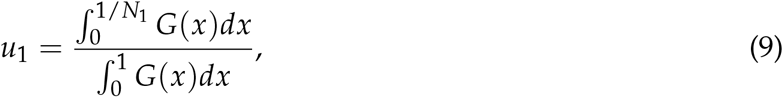

where 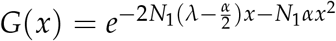.

Next, we derive the establishment probability of allele A that newly arises in subpopulation II with initial frequency 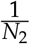. Because allele A is maladaptive in subpopulation II, we can assume that the frequency of allele A in subpopulation II should be kept low due to divergent selection, so that the process should be described by the branching process. The selection coefficient against allele A is 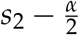 in subpopulation II. Then, the establishment probability of allele A which arises in subpopulation II is approximated by

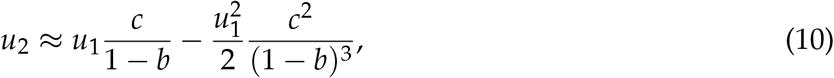

where 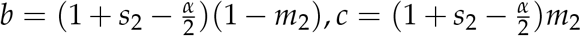. For the details of the derivation, see APPENDIX A.

The accuracy of our derivation was checked by simulations (Figure 1). Forward simulations were carried out under the haploid model to obtain *u*_1_ and *u*_2_, with initial frequency of allele A (*p*_1_, *p*_2_) = (1/*N*_1_, 0) and (0, 1/*N*_2_), respectively. For each parameter set, we ran ≥100,000 replications and counted the number of replications in which A establishes in the simulated population. The “establishment” is defined such that the new introduced mutation is still present after 5*N*_1_ generations passed. It should be noted that, according to our definition, the established replications include two cases; case C where alleles A and a coexist, namely, allele A is nearly fixed in subpopulation I while allele a is nearly fixed in subpopulation II, typical to strong divergent selection, and case F where allele A is fixed in both subpopulations. Let *Pc* be the relative proportion of case C in established replications. In Figure 1, a gray region is placed such that *Pc* > 0.9 in the left, while *Pc* < 0.1 in the right. According to our derivation, *u*_1_ and *u*_2_ from Equations 9 and 10 evaluate the sum of these two cases, so that they are comparable to the result of our simulations, although our interest is in case C where strong divergent selection is maintaining the two alleles (i.e., left of the gray region).

**Figure 1.**
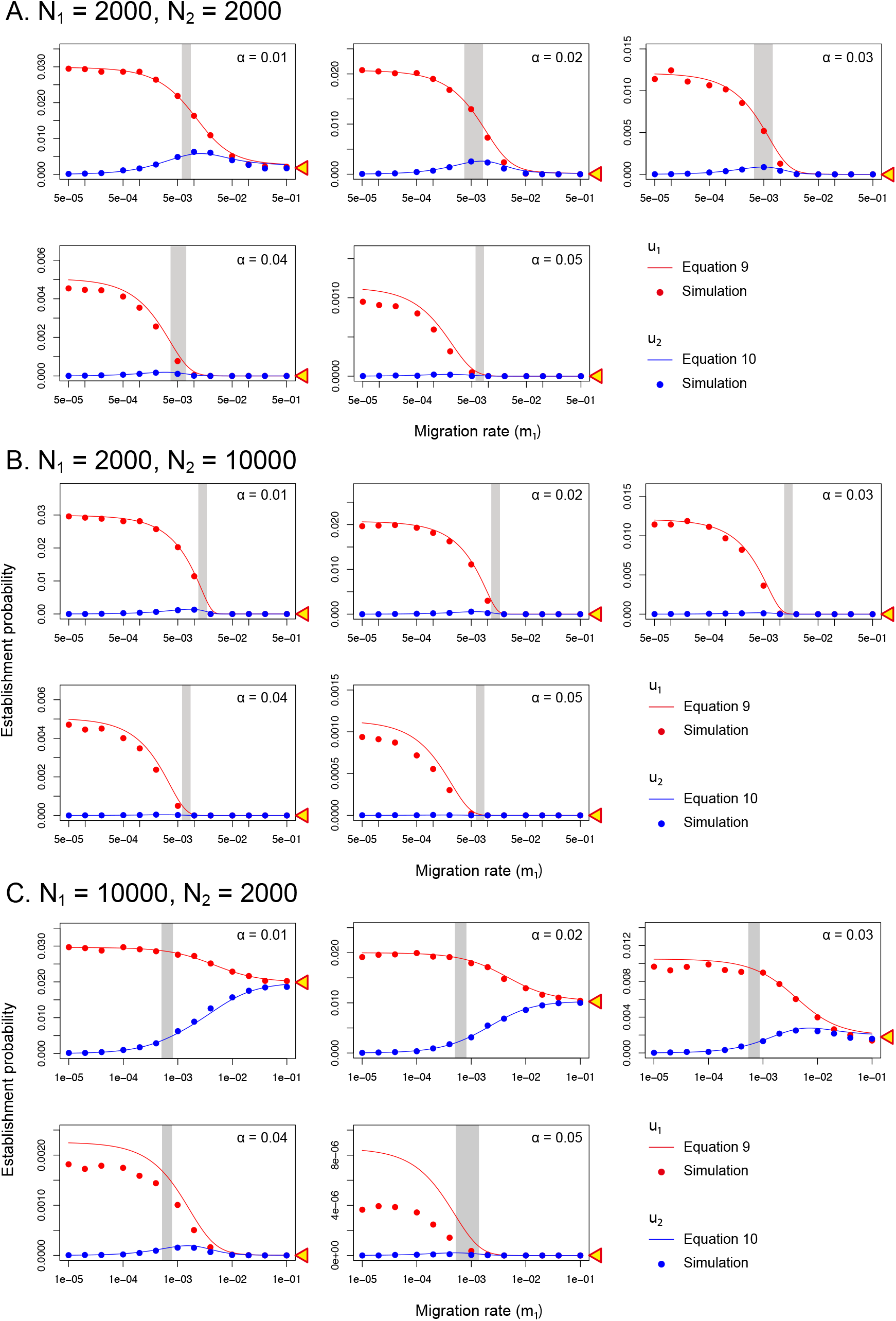
Establishment probability of a magic trait allele in the haploid model, plotted against the migration rate. *s*_1_ = 0.02 and *s*_2_ = −0.01 are fixed, and three pairs of the population sizes are assumed: (A) *N*_1_ = *N*_2_ = 2000, (B) *N*_1_ = 2000 and *N*_2_ = 10000, (C) *N*_1_ = 10000 and *N*_2_ = 2000. For each pair of population sizes, the strength of assortative mating is changed from *α* = 0.01 to 0.05. In each panel, a gray region is presented such that the proportion of the replications where two alleles (A and a) coexisted (*Pc*) > 0.9 in the left, while *Pc* < 0.1 in the right. The yellow triangle on the right vertical axis indicates the establishment probability assuming a very large migration rate.

Figure 1 show the establishment probability derived from Equations 9 and 10 as a function of migration rate, together with the simulation results. We fix *s*_1_ = 0.02 and *s*_2_ = −0.01, and the strength of assortative mating is changed (*α* = 0.01, 0.02, 0.03, 0.04, 0.05). In Figure 1, three pairs of *N*_1_ and *N*_2_ were considered; (A): *N*_1_ = *N*_2_ = 2000; (B): *N*_1_ = 2000, *N*_2_ = 10000; (C): *N*_1_ = 10000, *N*_2_ = 2000. It is found that Equations 9 and 10 generally agree well with the simulation result when the selection intensity is on the order of 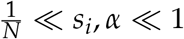.

When the effect of assortative mating is very weak (i.e., *α* = 0.01, the top left panel in Figures 1A, B, C), our results are overall similar to our previous work (Sakamoto and Innan 2019): when the migration rate is very low, the two subpopulations can be treated independently, so that *u*_1_ ~ 2*λ* where *λ* = *s*_1_ if we ignore assortative mating, while *u*_2_ ~ 0. As the migration rate increases, *u*_1_ decreases and *u*_2_ increases, and they become similar to each other for a large migration rate. Because allele A is advantageous only in subpopulation I, this beneficial effect would be reduced with increasing the migration rate, and vise versa for allele a. When the migration rate is very large (*m* ~ 0.5), the two subpopulations can be considered as a single random-mating population, and the fixation probability of a single mutation is mainly determined by the average selection coefficient,

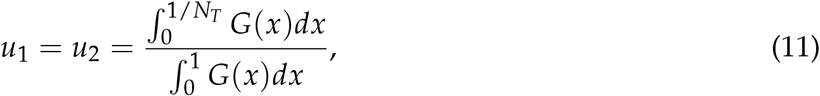

where 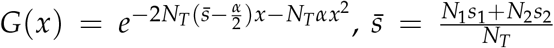 and *N*_*T*_ = *N*_1_ + *N*_2_ (presented by a triangle in Figure 1). The gray region for 0.1 ≤ *Pc* ≤ 0.9 is placed in a narrow window of the migration rate, indicating that a fairly small difference in the migration rate around the gray region could change the typical outcome dramatically. In the right of the gray region, the established probability essentially means that allele A is fixed in both subpopulations, whereas in the left, the two alleles are nearly fixed in each subpopulation (i.e., divergent selection is maintaining them).

In the following, the effect of *α* on *u_i_* is explained along with the symmetric case (*N*_1_ = *N*_2_ = 2000) shown in Figure 1A. Allele A is favored in subpopulation I through divergent selection, which is a positive force for its establishment, although this positive effect is weakened by migration. Mathematically, *λ* in Equation 7 can be considered as the effective intensity of divergent selection, where migration is taken into account. In contrast, assortative mating is a negative force when allele A is rare, which works to make allele A difficult to increase in frequency. This is because allele A has to mate with allele a in most cases (i.e., reproduction is less successful). Thus, the relative contributions of divergent selection and assortative mating largely determines the fate of allele A.

Let us first assume a very small migration rate (see the left end at *m*_1_ = 5 × 10 ^−5^ in each panel in Figure 1A). If we ignore migration, *λ* is simply given by *s_i_* and the total strength of selection (divergent selection and assortative mating) on allele A is roughly given by 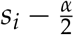 in subpopulation *i*. As expected, the established probability *u*_1_ and *u*_2_ decrease with increasing *α*. When assortative mating is weak (*α* = 0.01 ~ 0.03 in Figure 1A), *u*_1_ could be roughly approximated by *u*_1_ = 2*s*_1_ − *α*. When *α* = 0.04, we have 2*s*_1_ − *α* = 0, where the two selective forces should roughly cancel each other, but it seems that *u*_1_ exceeds the neutral expectation (1/*N*_1_) because the negative effect of assortative mating is relaxed when *p*_1_ increases. The establishment probability *u*_1_ is very low for *α* > 0.04, where selection in total works against allele A due to strong assortative mating. It should be interesting to point out that, even with strong assortative mating, allele A can establish if A happens to increase in frequency by random genetic drift. Once allele A becomes common, the negative effect of assortative mating is somehow reduced, and it could successfully go to the establishment by the positive effect of divergent selection.

As the migration rate increases, the intensity of divergent selection is effectively weaken, so that *u*_1_ decreases. At the limit of free migration, we have *u*_1_ = *u*_2_ is given by Equation 11 as mentioned above. The location of the gray region is quite constant over the range of *α*. This robustness to *α* should be because the eventual fate of A (i.e., cases C or F) is mainly determined by divergent selection because assortative mating works efficiently only when allele A is rare, namely, shortly after it arises in the population. Once allele A increases and exceed a certain frequency, allele A may not be so deleterious because there are many allele A to mate with no reproductive reduction.

Focusing on *u*_1_, Figure 2 explores a wider parametric space for *s*_1_ and *α* while the other parameters are fixed (*m*_1_ = *m*_2_ = 0.005 and *s*_2_ = −0.02). To demonstrate the effect of the population size, three population sizes are considered assuming the same population sizes in the two subpopulations (*N* = *N*_1_ = *N*_2_ = 2000, 6000, 20000). In each panel, analytical and simulation results are shown: Analytical results from Equation 2 are presented by the color of the background, while colors inside the circles represent simulation results.

**Figure 2.**
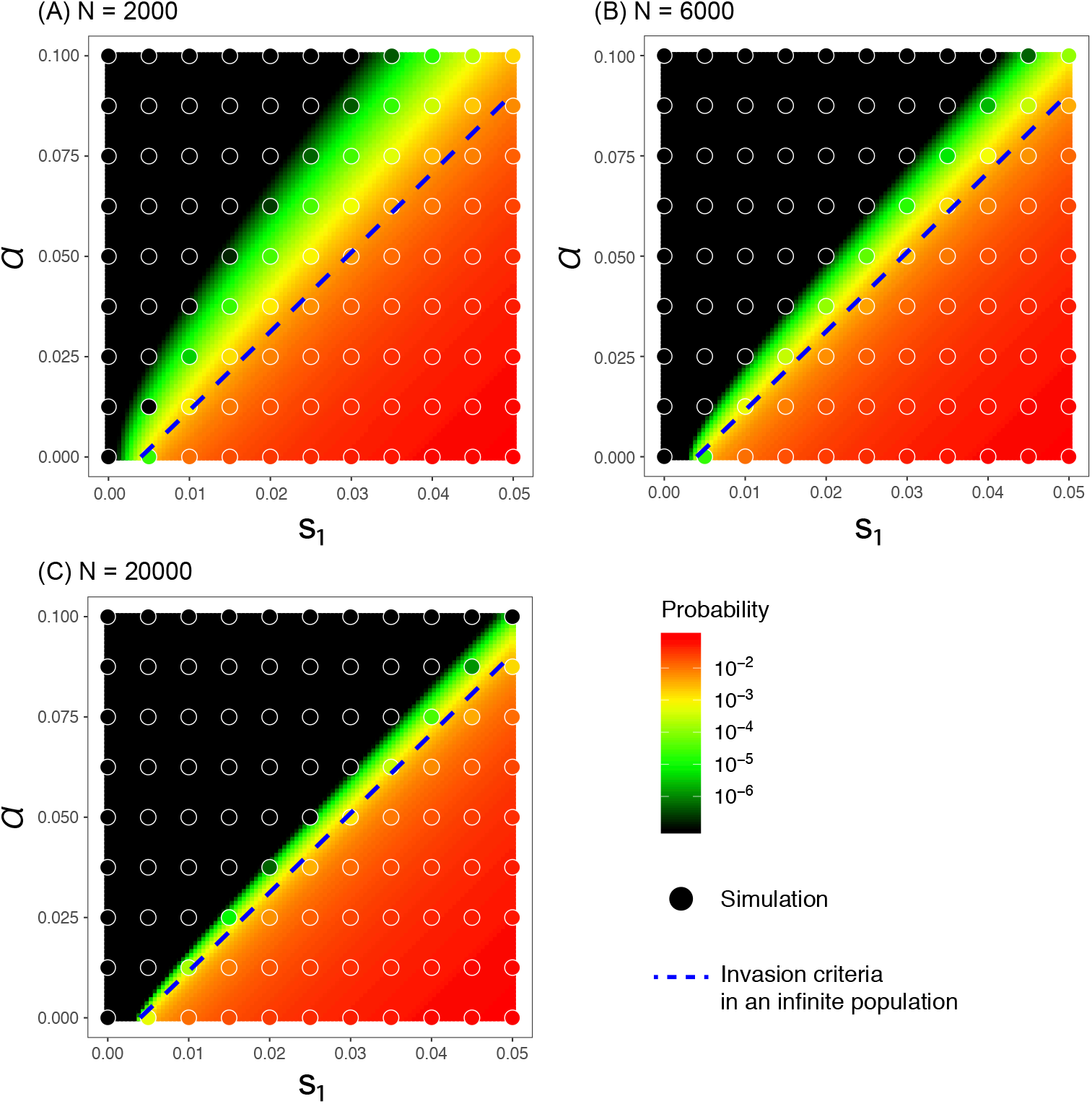
Establishment probability of a magic trait allele in the haploid model with different population sizes (*N*_1_ = *N*_2_ = 2000, 6000, 20000). *m*_1_ = *m*_2_ = 0.005 and *s*_2_ = −0.02 are fixed. The background color presents the theoretical approximation (Equation 9) while circle’s color presents the simulation result. The blue dashed line presents the invasion criteria assuming an infinite population (see text for details).

Again, the overall agreement of colors in the circle and those in the background suggest excellent performance of Equation 9 for a wide range of the parameter set. It seems that our analytical result slightly overestimates the establishment probability when *s_i_* and *α* are so large that the establishment probability is sensitive to the second order of *s_i_*, *α*, which were ignored in our derivation. The blue dashed line represents 2*λ* = *α*, where the two selective forces roughly cancel each other so that it can be theoretically considered as a threshold of establishment in an infinite population. When the population size is very large, *u*_1_ drops quickly above the line, whereas allele A may establish even above the line when the population size is small, because of random genetic drift.

### Establishment Probability in the Diploid Model

The initial state is that all individuals are aa homozygotes in both subpopulations. We first consider the establishment probability of allele A which newly arises in subpopulation I with initial frequency 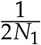. As well as our derivation for the haploid model, we approximate the two-population process by a one-population system by focusing the establishment process in subpopulation I alone (Yeaman and Otto 2011). To do so, the fitnesses of genotypes AA, Aa and aa are given by 1 + *λ*, 1 + *hλ* and 1, respectively, where *λ* is the effective strength of natural selection as defined above. Allele A can be considered dominant over allele a when *h* = 1, and recessive when *h* = 0. Following Yeaman and Otto (2011), we assume that the leading eigenvalue of transition matrix approximates the growth rate of allele A in a virtual one-population system (when allele A is rare). Then, *λ* is given by

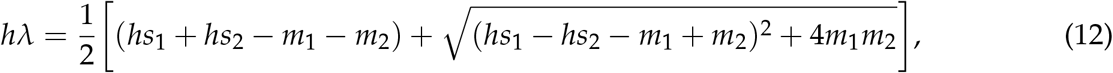

where we assume fairly strong selection, say *hs*_*i*_ is as large as on the order of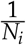.

Given Equation 12, *M*_*p*_ and *M_q_*, the expected changes in the genotype frequencies of AA and Aa in one generation, are obtained by assuming that *λ*, *α* ≪ 1 and ignoring the second order of these, and the Kolmogorov’s forward equation is given by

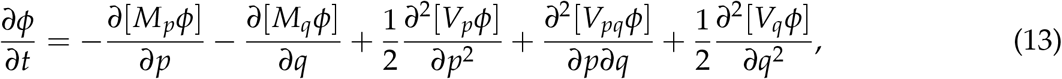

where 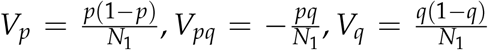 and *ϕ* is an arbitrary transition density. By changing the variables, Equation 13 is rewritten as a function of allele frequency of A, *x*, and frequency of heterozygote *q* as

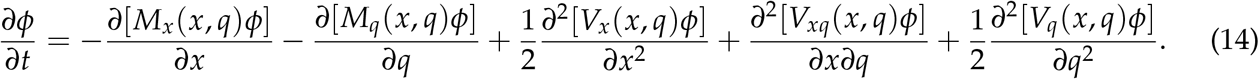

This forward equation with two variables, *x* and *q*, is still difficult to solve, so that we attempt to reduce the dimension with approximation, that is, the frequency of heterozygotes, *q*, is approximated by a function of *x*. If we ignore assortative mating so that the Hardy-Weinberg equilibrium holds, we can simply assume *q*(*x*) = 2*x*(1 − *x*). Even with assortative mating, Yamamichi and Sasaki (2013) demonstrated that the assumption of the Hardy-Weinberg equilibrium (i.e., *q*(*x*) = 2*x*(1 − *x*)) works fairly well. Recently, Newberry *et al.* (2016) proposed that *q*(*x*) can be given by the solution of the differential equation 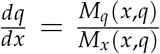 which satisfies the condition lim_*x*→0_ *q*(*x*) = 0. We here employ the method of Newberry *et al.* (2016) and obtain *q*(*x*) ≈ 2*x*(1 − *x*) − 4*x*^2^(1 − *x*)^2^(*x*^2^ − 2*hx* + *h*)*α* by a perturbation approach (for details see APPENDIX B). With this approximation of *q*(*x*) and by ignoring the stochastic deviation of the frequency of heterozygote from *q*(*x*), Equation 14 can be reduced to a one-dimensional diffusion equation:

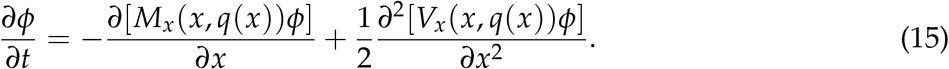

Then, by using Kimura’s formula (Kimura 1962), the establishment probability of allele A that newly arises in subpopulation I, *u*_1_, is given by

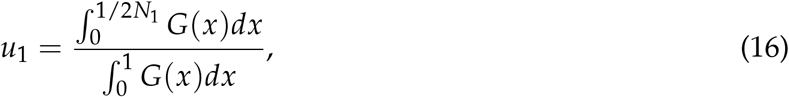

where 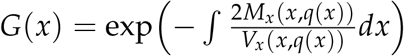.

Next, we derive the establishment probability of allele A that newly arises in subpopulation II. Following the haploid case, the establishment probability *u*_2_ is given by

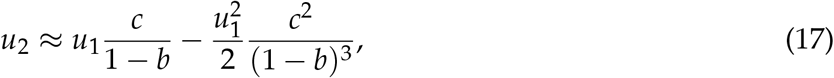

where 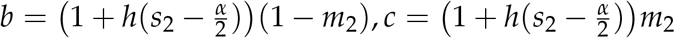.

The establishment probability derived from Equations 16 and 17 is compared with simulation in Figure 3. Simulations were performed by assuming *N* = *N*_1_ = *N*_2_ = 1000, *s*_1_ = 0.02, *s*_2_ = −0.01 and *α* was changed from 0.01 to 0.05. Three different degrees of dominance were considered; complete dominance (*h* = 1), additive (*h* = 0.5), and nearly recessive (*h* = 0.05). Overall, our analytical results again agree well with the simulation result when 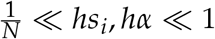. We obtain almost the same result as the haploid model when *h* = 1 (Figure 3A), and *u*_1_ and *u*_2_ decrease as *h* decreases (Figures 3B and C). This trend can be explained if we consider how selection works on allele A in the very early phases, namely, when the frequency is very low. When *h* = 1, a newly arisen allele A exhibit a full phenotype difference, which is immediately subject to selection. This is a similar situation to the haploid model. For *h* < 1, the phenotypic effect of allele A could be reduced to some extent (i.e., by a factor of *h*), therefore the selective pressure could be relaxed. Thus, the selective pressure on allele A in the diploid model can be summarized by *hs*_*i*_ and *hα* in the early phases. As expected, if we plot *u*_1_ for different *h* where the horizontal and vertical axises are adjusted by *hs*_*i*_ and *hα*, respectively, the distribution of the establishment probability are very similar (Figure 4).

**Figure 3.**
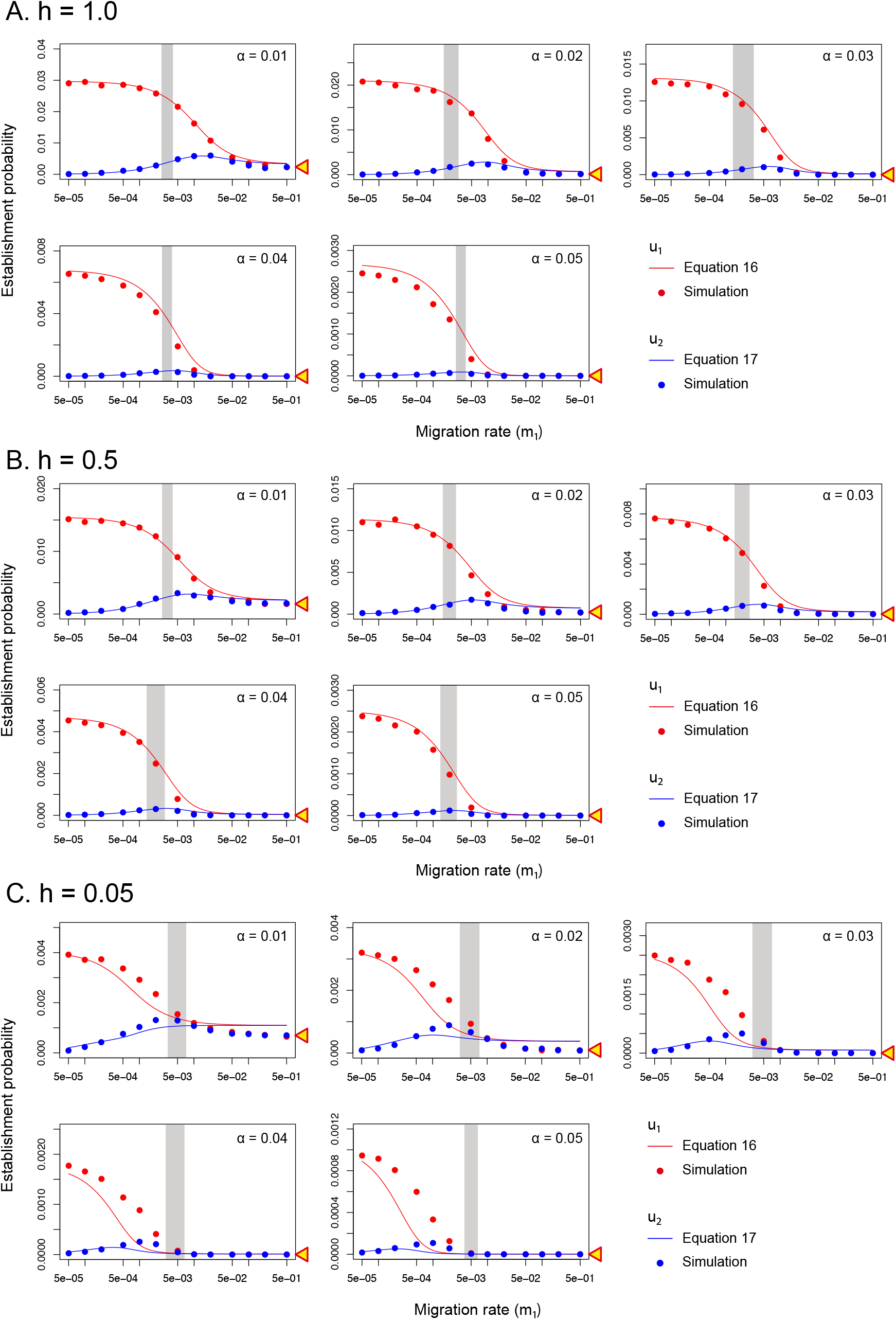
Establishment probability of a magic trait allele in the diploid model, plotted against the migration rate. *s*_1_ = 0.02, *s*_2_ = −0.01 and *N*_1_ = *N*_2_ = 1000 are fixed, and three dominance coefficients are assumed: (A) *h* = 1.0, (B) *h* = 0.5, (C) *h* = 0.05. For each dominance coefficient, the strength of assortative mating is changed from *α* = 0.01 to In each panel, a gray region is presented such that the proportion of the replications where two alleles (A and a) coexisted (*Pc*) > 0.9 in the left, while *Pc* < 0.1 in the right. The yellow triangle on the right vertical axis indicates the establishment probability assuming a very large migration rate.

**Figure 4.**
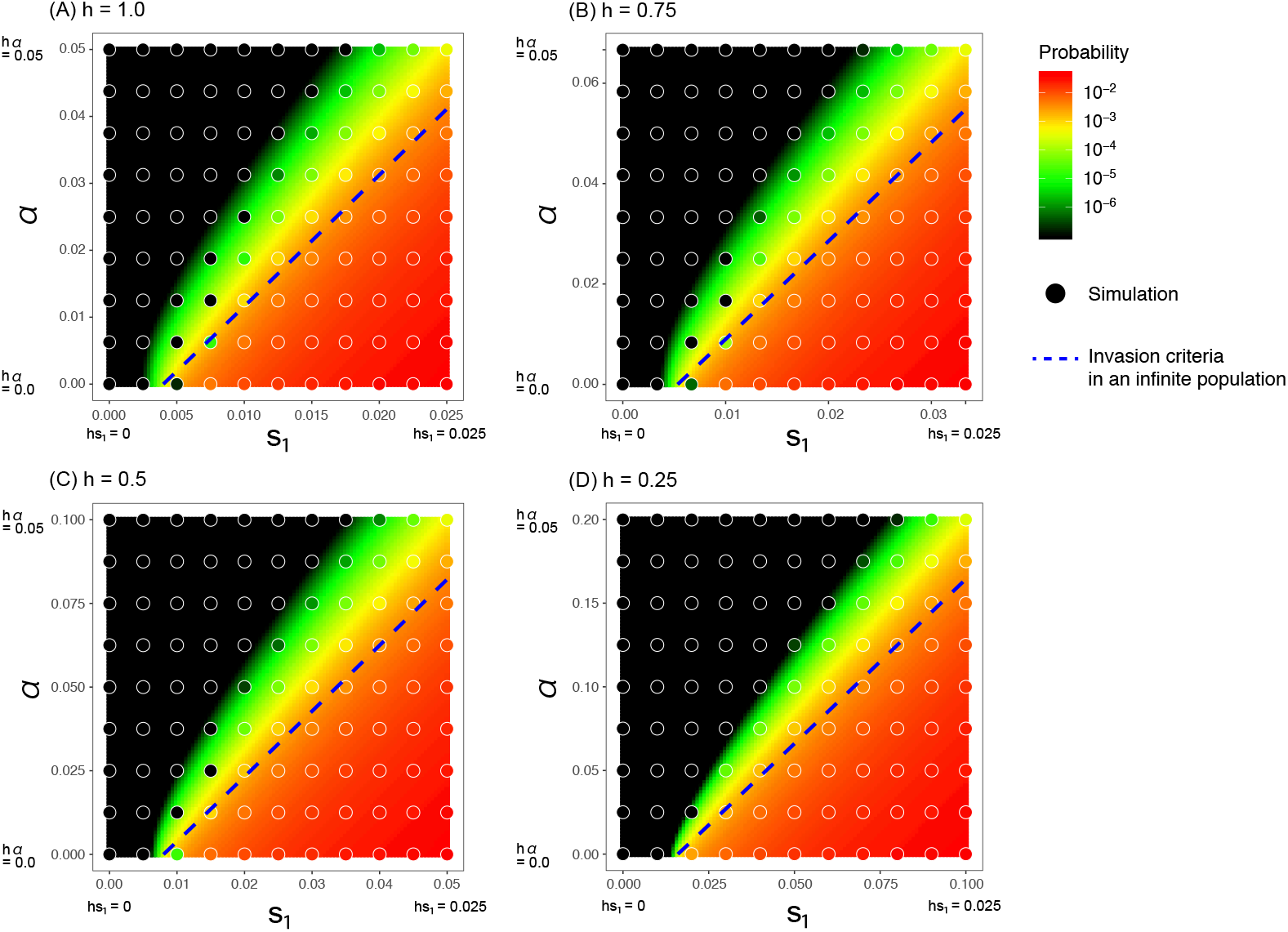
Establishment probability of a magic trait allele in the diploid model in different level of dominance (*h* = 1.0, 0.75, 0.5, 0.25). *hs*_2_ = −0.02 is fixed. The background color presents the theoretical approximation (Equation 16) while circle’s color presents the simulation result. The blue dashed line presents the invasion criteria assuming an infinite population (see text for details). The results are plotted such that the horizontal and vertical axises are scaled by *hs*_1_ and *hα*, respectively, therefore the blue dashed lines show up at the same position in each panel.

### Establishment Trajectory of Allele Frequency

When allele A establishes, it rapidly spreads in the subpopulation I and is stably maintained at frequencies around the migration-selection balance, 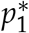. We consider the frequency trajectory of allele A under the condition that it establishes. In practice, we obtain the mean sojourn time of allele A until it reaches frequency 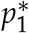 for the first time under the condition where allele A reaches frequency 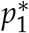.

In the haploid model, assuming a low migration rate and strong selection, 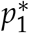 is given by 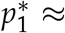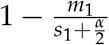. Following Ewens (1973), we can derive the conditional mean sojourn time at frequency *x*, *T*\*(*x*) as

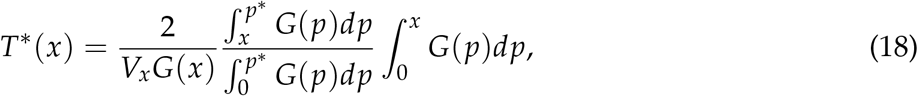

where 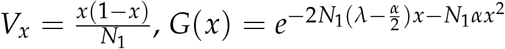. We then obtain the time required for a newly arisen allele A to reach allele frequency *x*:

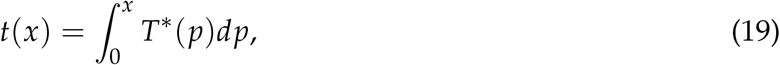

so that 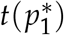 provides the establishment time. Note that although this argument holds for a new allele arisen in subpopulation I, it can be applied to an allele arisen in subpopulation II after it migrated into subpopulation I.

In the diploid model, we can approximate 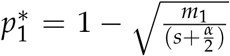 for *h* ~ 1 and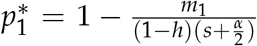 for other cases *h* is relatively smaller than 1. We then obtain the conditional mean sojourn time at frequency *x* by using Equation 18 by replacing *V_x_* = *V_x_* (*x*, *q*(*x*)) and 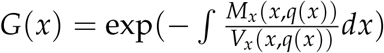. Our derivation works well when *h* is not small.

The theoretical result is quite simple and there is no marked difference between the haploid and diploid models. Therefore, we demonstrate the point by using the haploid model. Figure 5A shows the theoretical trajectory of *t*(*x*) by the red line when strong assortative mating is working (*α* = 0.04), compared with the case with no assortative mating (*α* = 0, Figure 5B). For each panel, we also show ten independent established runs by black lines. The major difference is in that the allele frequency likely stays long in low frequency with assortative mating (Figure 5A), in comparison with the symmetric function in Figure 5B. This difference is easy to understand: As mentioned above, the negative effect on the newly arisen allele is strong only when its allele frequency is low, and the effect is relaxed once it increases. This pattern is globally observed both in the haploid and diploid models.

**Figure 5.**
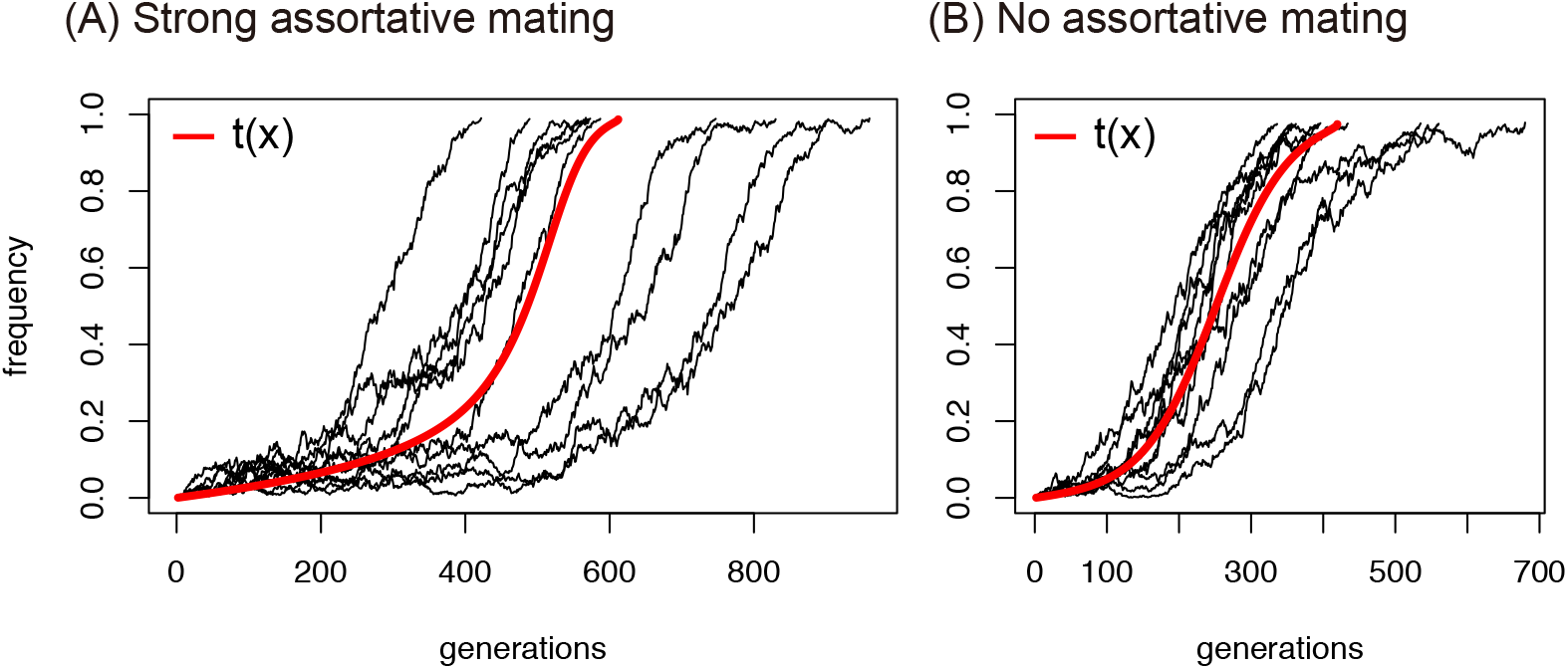
Trajectory of the frequency of allele A in the haploid model. *s*_1_ = 0.02, *s*_2_ = −0.01, *m*_1_ = *m*_2_ = 0.0005 and *N*_1_ = *N*_2_ = 2000 are assumed. (A) with strong assortative mating *α* = 0.04 and (B) with no assortative mating *α* = 0. The black lines shows 10 independent paths from simulations and the red line is the theoretical prediction.

## DISCUSSION

We are interested in how a newly arisen allele of a magic trait (allele A) behaves and contributes to ecological speciation. A magic trait is defined such that a single trait is subject to both divergent selection and assortative mating. Divergent selection simply favors the new allele to fix where it is beneficial, thereby creating a genetic difference between subpopulations. Assortative mating works in a more trick way: it also prefers difference between subpopulations but a new allele is not always advantageous. This is because when allele A is still rare after it arises, allele A is disadvantageous because allele A has to mate with allele a in most cases with a reduction in reproductive success. Once allele A becomes common so that A and A can mate with no reduction in reproduction, allele A is not very deleterious any more and would fix in the adapted subpopulation, thereby contributing to genetic divergence between subpopulations.

In this work, we investigate the establishment probability of such a magic trait allele under the haploid and diploid models. We successfully obtained the establishment probability by using the approximation method of Yeaman and Otto (2011). We confirmed that our derivation agreed well with our simulation results. Our theory mainly focuses on the early phases, that is, when allele A is still rare so that divergent selection and assortative mating counteract. It is theoretically demonstrated that the relative contributions of divergent selection and assortative mating largely determine the fate of allele A.

In the haploid model, *λ* in Equation 7 explains the effective intensity of divergent selection with migration taken into account, and the intensity of assortative mating is parameterized by *α*. Theoretically, 2*λ* − *α* is the key quantity to determine the fate of allele A when its frequency (*p*_1_) is rare. If 2*λ* − *α* > 0, allele A is on average selected for, and selected against if 2*λ* − *α* < 0 in an infinite population. In a finite population, we show that allele A can establish even when 2*λ* − *α* < 0. This is especially true for a small population (Figures 2, S1-S3) because random genetic drift occasionally increase *p*_1_, and the negative effect of assortative mating is relaxed once allele A becomes common. Then, it is likely that allele A goes to establishment.

The results for the diploid model is very similar to that for the haploid model. The major difference is that the dominance effect is involved in the diploid model. It is demonstrated that the establishment process can be well described if both *s_i_* and *α* are scaled by *h*.

We also explore how the allele frequency behaves along the establishment. We theoretically obtained the trajectory of allele frequency, which clearly demonstrated that the negative effect of assortative mating retards establishment while this effect is relaxed once the allele frequency increases.

In summary, our theory well demonstrates the behavior of a magic trait allele that are subject to divergent selection and assortative mating. We also show the establishment of a magic trait allele is largely affected by the population size, because the fate of a newly arisen allele is mainly determined when it is rare, where random genetic drift plays a central role. The theoretical results will enhance our understanding of how natural selection initiates speciation.

## APPENDIX

## APPENDIX A: Establishment probability of allele A that arises in subpopulation II

We here derive the establishment probability of a single allele A which arises in the maladapted subpopulation II. First, we derive the probability distribution of the number of migrant alleles to subpopulation I which originate from a single allele A in subpopulation II by using the branching process. Then, by assuming that migrant alleles behave independently, we derive an approximate formula of the establishment probability. The derivation works for both haploid and diploid models.

We put the probability variables *X*_*t*_ as the number of allele A in subpopulation II at generation *t* and *Y*_*t*_ as the number of allele A which have migrated to subpopulation I until generation *t*. Because we consider the case where negative selection works on allele A in subpopulation II, we can assume that the frequency of allele A is kept low so that allele A mostly exist in heterozygotes in the diploid case. Therefore, *X*_*t*_ is almost same as the number of heterozygotes in the diploid model. We define a probability generating function at generation *t* as 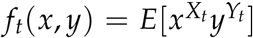. We set the generating function of joint distribution of the number of allele A in subpopulation II and the number of allele A that have just migrated to subpopulation I which originates from one allele A at the previous generation as *h*(*x*, *y*). If we assume the number of offsprings follows the poisson distribution, 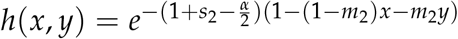 in the haploid case. Then, we have

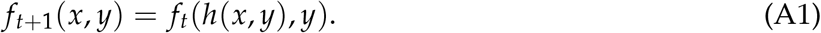

We put 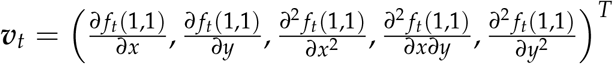. By using Equation A1, we obatin

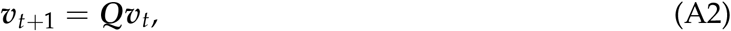

where

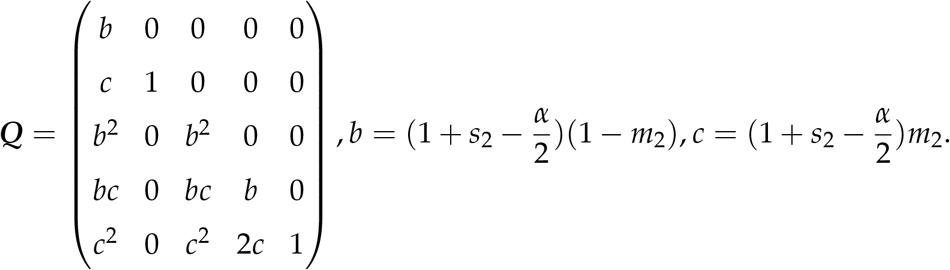

When *t* goes to infinity, 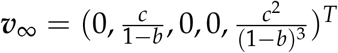.

Then, the establishment probability of a single allele A that arises in the maladapted subpopulation II, *u*_2_, is approximately given by

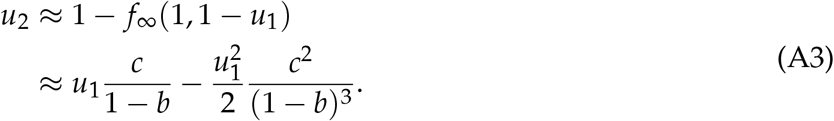

For the diploid case, we should substitute *s*_2_ and *α* by *hs*_2_ and *hα* in the above equations.

## APPENDIX B: Approximation of the frequency of heterozygotes

We derive an approximate formula which describes the frequency of heterozygotes in the diploid model. We assume that the strength of assortment mating, *α*, is small and the frequency of heterozygote *q*(*x*) can be expanded as

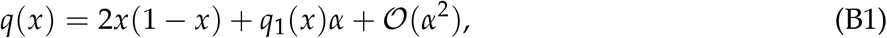

where *x* is the frequency of allele A. Following Newberry *et al.* (2016), we assume that *q*(*x*) is the solution of the differential equation, 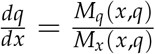. By substituting Equation B1, we have

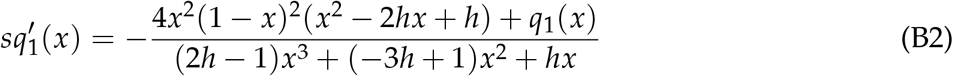

Because we assume *s* ≪ 1, we can derive *q*_1_(*x*) ≈ −4*x*^2^(1 − *x*)^2^(*x*^2^ − 2*hx* + *h*) by ignoring the left side of Equation B2.

---

**Figure S1.**
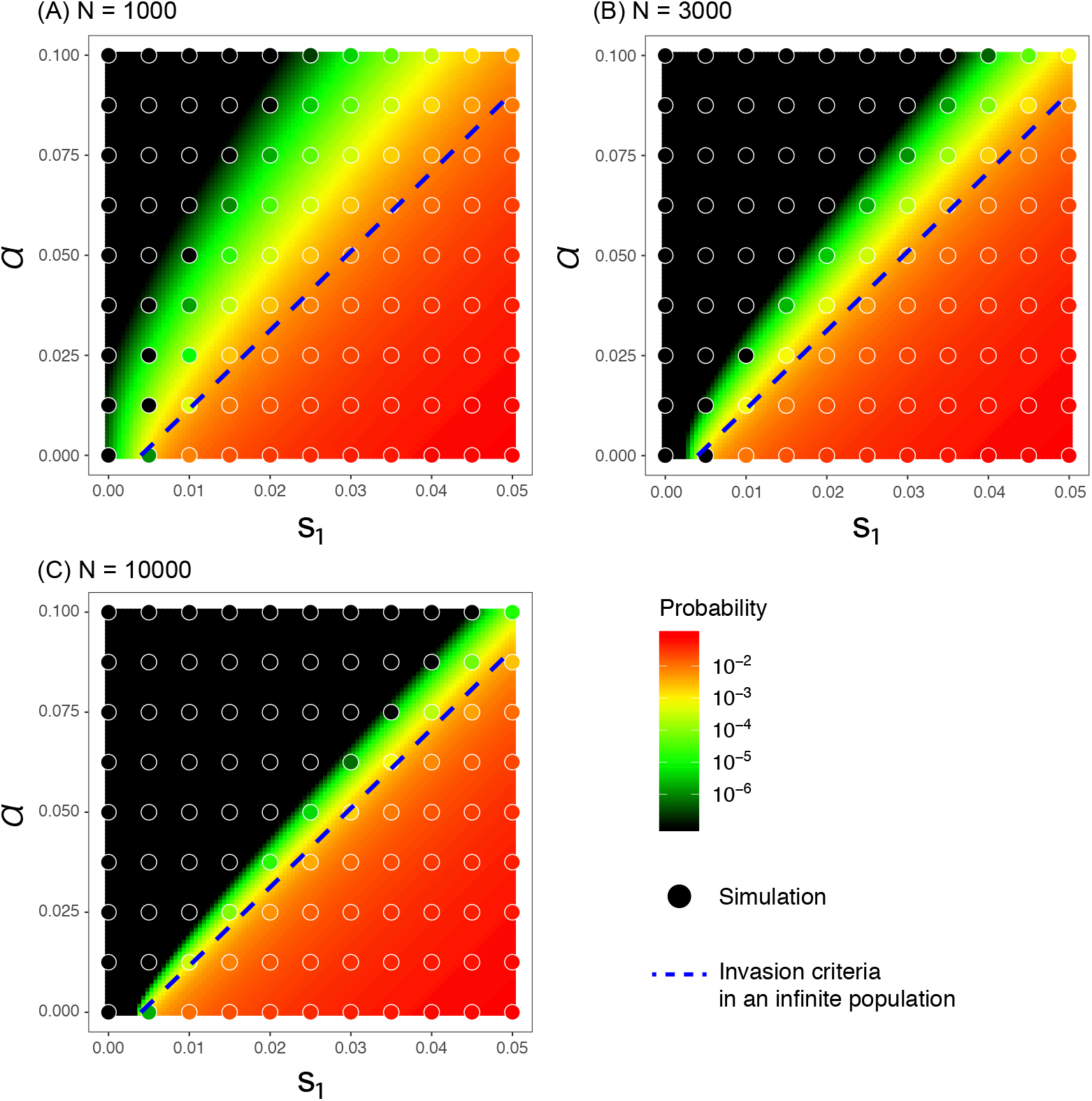
Establishment probability of a magic trait allele in the complete dominant case (*h* = 1.0) for different population size (*N*_1_ = *N*_2_ = 1000, 3000, 10000). *m*_1_ = *m*_2_ = 0.005 and *s*_2_ = −0.02 are assumed. The background color presents the theoretical approximation (Equation 16) while circle’s color presents the simulation result.

**Figure S2.**
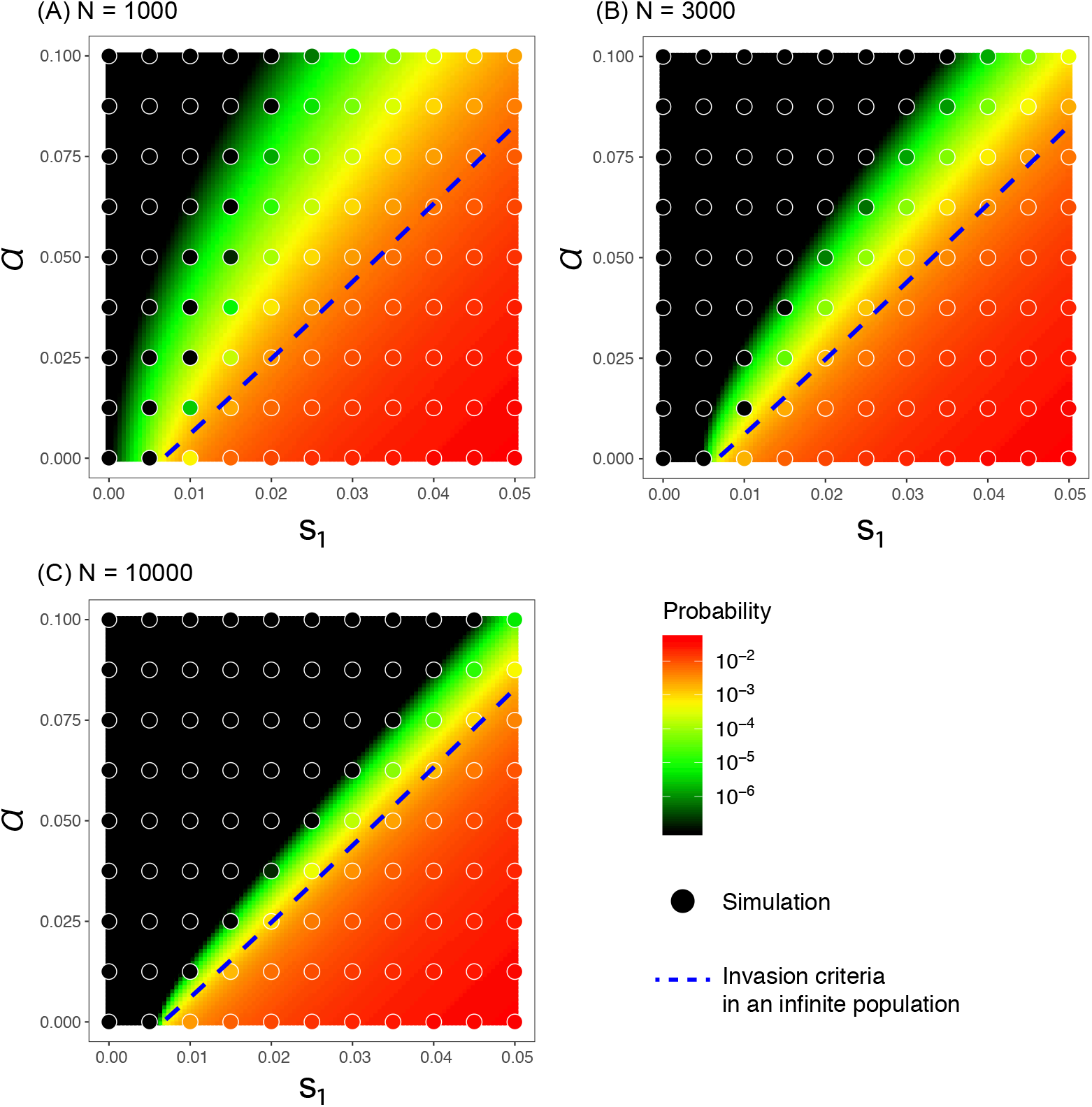
Establishment probability of a magic trait allele in the additive case (*h* = 0.5) for different population sizes (*N*_1_ = *N*_2_ = 1000, 3000, 10000). Other parameters are the same as figure S1.

**Figure S3.**
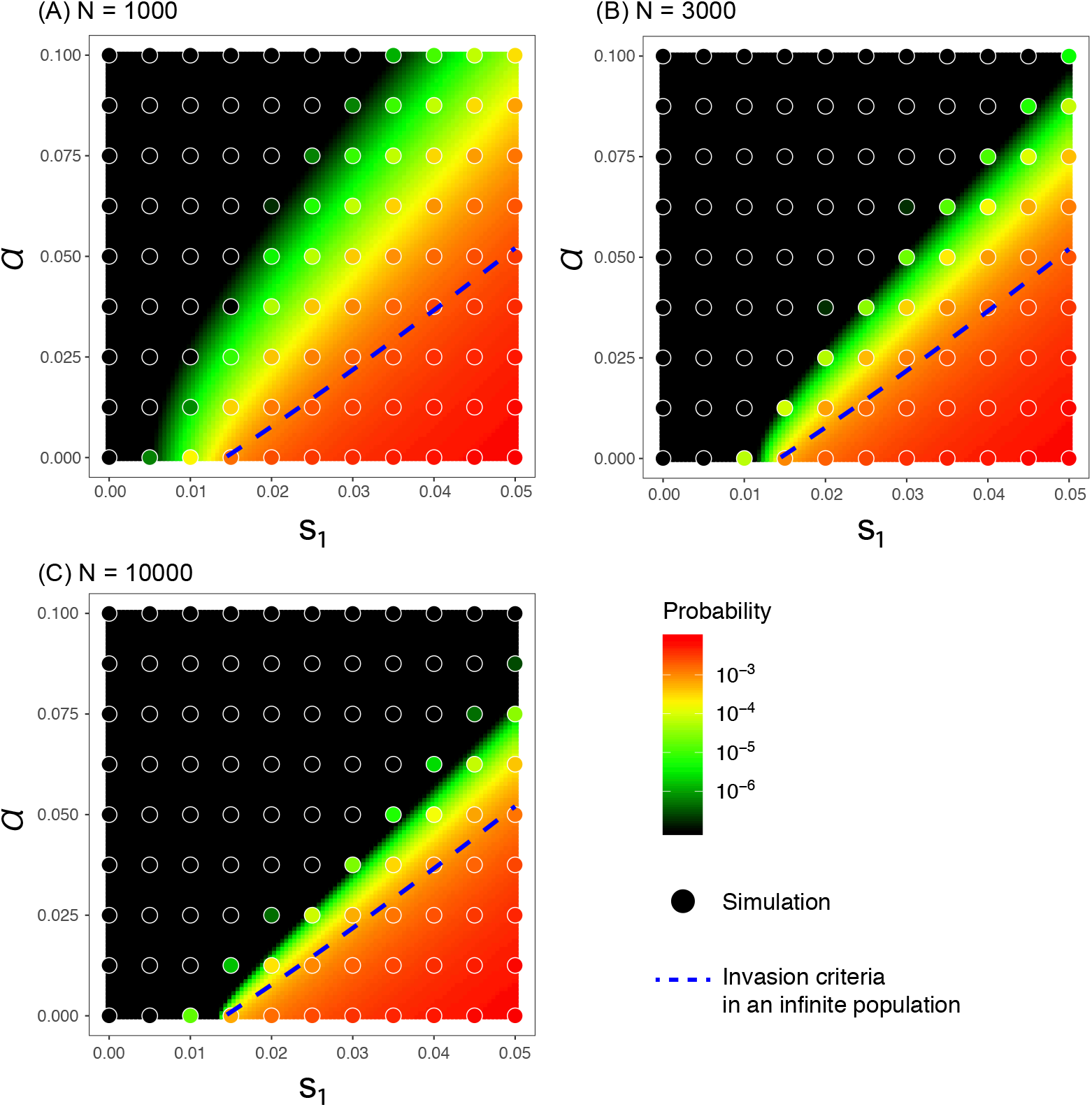
Establishment probability of a magic trait allele in a nearly recessive case (*h* = 0.05) for different population sizes (*N*_1_ = *N*_2_ = 1000, 3000, 10000). Other parameters are the same as figure S1.

## Literature Cited

Arnegard, M. E. and A. S. Kondrashov, 2004 Sympatric speciation by sexual selection alone is unlikely. Evolution 58: 222–237.

Barton, N. H., 1987 The probability of establishment of an advantageous mutant in a subdivided population. Genet. Res. 50: 35–40.

Bürger, R. and K. A. Schneider, 2006 Intraspecific competitive divergence and convergence under assortative mating. Am. Nat. 167: 190–205.

Cotto, O. and M. R. Servedio, 2017 The roles of sexual and viability selection in the evolution of incomplete reproductive isolation: From allopatry to sympatry. Am. Nat. 190: 680–693.

Coyne, J. A. and H. A. Orr, 2004 Speciation. Sinauer.

Dieckmann, U. and M. Doebeli, 1999 On the origin of species by sympatric speciation. Nature 400: 354–357.

Doebeli, M. and U. Dieckmann, 2003 Speciation along environmental gradients. Nature 421: 259–264.

Ewens, W., 1973 Conditional diffusion processes in population genetics. Theor. Popul. Biol. 4: 21–30.

Gavrilets, S., 2004 Fitness landscapes and the origin of species. Princeton University Press.

Higashi, M., G. Takimoto, and N. Yamamura, 1999 Sympatric speciation by sexual selection. Nature 402: 523–526.

Kimura, M., 1962 On the probability of fixation of mutant genes in a population. Genetics 47: 713–719.

Kirkpatrick, M. and S. L. Nuismer, 2004 Sexual selection can constrain sympatric speciation. Proc. R. Soc. Lond. B 271: 687–693.

Kirkpatrick, M. and V. Ravigné, 2002 Speciation by natural and sexual selection: models and experiments. Am. Nat. 159: S22–S35.

Kisdi, É. and T. Priklopil, 2011 Evolutionary branching of a magic trait. J. Math. Biol. 63: 361–397.

Kopp, M., M. R. Servedio, T. C. Mendelson, R. J. Safran, R. L. Rodríguez, M. E. Hauber, E. C. Scordato, L. B. Symes, C. N. Balakrishnan, D. M. Zonana, et al., 2018 Mechanisms of assortative mating in speciation with gene flow: connecting theory and empirical research. Am. Nat. 191: 1–20.

Maan, M. E. and O. Seehausen, 2011 Ecology, sexual selection and speciation. Ecol. Lett. 14: 591–602.

Matessi, C., A. Gimelfarb, and S. Gavrilets, 2001 Long-term buildup of reproductive isolation promoted by disruptive selection: how far does it go? Selection 2: 41–64.

Newberry, M. G., D. M. McCandlish, and J. B. Plotkin, 2016 Assortative mating can impede or facilitate fixation of underdominant alleles. Theor. Popul. Biol. 112: 14–21.

Otto, S. P., M. R. Servedio, and S. L. Nuismer, 2008 Frequency-dependent selection and the evolution of assortative mating. Genetics 179: 2091–2112.

Pennings, P. S., M. Kopp, G. Meszéna, U. Dieckmann, and J. Hermisson, 2008 An analytically tractable model for competitive speciation. Am. Nat. 171: E44–E71.

Pollak, E., 1966 On the survival of a gene in a subdivided population. J. Appl. Prob. 3: 142–155.

Rettelbach, A., M. Kopp, U. Dieckmann, and J. Hermisson, 2013 Three modes of adaptive speciation in spatially structured populations. Am. Nat. 182: E215–E234.

Ritchie, M. G., 2007 Sexual selection and speciation. Annu. Rev. Ecol. Evol. Syst. 38: 79–102.

Sakamoto, T. and H. Innan, 2019 The evolutionary dynamics of a genetic barrier to gene flow: From the establishment to the emergence of a peak of divergence. Genetics 212: 1383–1398.

Servedio, M. R. and J. W. Boughman, 2017 The role of sexual selection in local adaptation and speciation. Annu. Rev. Ecol. Evol. Syst. 48: 85–109.

Servedio, M. R. and R. Bürger, 2015 The effects of sexual selection on trait divergence in a peripheral population with gene flow. Evolution 69: 2648–2661.

Servedio, M. R., G. S. Van Doorn, M. Kopp, A. M. Frame, and P. Nosil, 2011 Magic traits in speciation:‘magic’but not rare? Trends Ecol. Evol. 26: 389–397.

Slatkin, M., 1982 Pleiotropy and parapatric speciation. Evolution 36: 263–270.

Takimoto, G., M. Higashi, and N. Yamamura, 2000 A deterministic genetic model for sympatric speciation by sexual selection. Evolution 54: 1870–1881.

Thibert-Plante, X. and S. Gavrilets, 2013 Evolution of mate choice and the so-called magic traits in ecological speciation. Ecol. Lett. 16: 1004–1013.

Tomasini, M. and S. Peischl, 2018 Establishment of locally adapted mutations under divergent selection. Genetics 209: 885–895.

Turner, G. F. and M. T. Burrows, 1995 A model of sympatric speciation by sexual selection. Proc. R. Soc. Lond. B 260: 287–292.

Weissing, F. J., P. Edelaar, and G. S. Van Doorn, 2011 Adaptive speciation theory: a conceptual review. Behav. Ecol. Sociobiol. 65: 461–480.

Wu, C.-I., 1985 A stochastic simulation study on speciation by sexual selection. Evolution 39: 66–82.

Yamamichi, M. and A. Sasaki, 2013 Single-gene speciation with pleiotropy: effects of allele dominance, population size, and delayed inheritance. Evolution 67: 2011–2023.

Yeaman, S. and S. P. Otto, 2011 Establishment and maintenance of adaptive genetic divergence under migration, selection, and drift. Evolution 65: 2123–2129.

